# DTPPI: predicting drug interactions using a weighted drug-protein network

**DOI:** 10.1101/2025.01.06.631638

**Authors:** Szymon Szydlik, Golnaz Taheri

## Abstract

Polypharmacy, the practice of using multiple drugs to treat complex diseases, poses a significant risk of drug-drug interactions (DDIs), which can lead to unanticipated adverse drug reactions (ADRs) and toxicity. Identifying and understanding these DDIs is crucial to ensuring the safety of polypharmacy. Traditional laboratory-based methods for detecting DDI are costly and time consuming, prompting the development of computational approaches. However, many of these methods face limitations, mainly the lack of utilization of biological networks to model drug mechanics. Such an approach could lead to a new technique with better and more accurate DDI predictions. In response to these challenges, we propose the DTPPI network, a novel machine learning approach that leverages a drug-target-protein-protein interaction network to improve DDI prediction. By extracting topological features and combining them with biological drug features, the DTPPI method enhances the performance of a multilayer perceptron model. The evaluation results showed an AUC of 0.64 for topological characteristics alone, 0.89 for biological characteristics, and 0.91 for combined features, demonstrating that integrating topological and biological data significantly improves the prediction accuracy of DDI. Materials and implementations are available at: https://github.com/Golnazthr/DTPPI

**Highlights:**

1. Constructs a weighted graph network to model interactions among drugs, proteins, and targets, applicable to all drug types.
2. Extracts six universal topological features from the graph, independent of chemical structure.
3. Enhances DDI prediction by using topological features alone or alongside traditional drug features.
4. Incorporates an MLP model for superior predictive accuracy using combined features.

## 1. Introduction

Polypharmacy, also known as polytherapy, refers to a treatment approach that involves the administration of two or more medications simultaneously to enhance the effectiveness of treatment. This method is often used to address multiple symptoms or conditions at once [1, 2]. Polypharmacy is particularly common among elderly patients with multiple chronic illnesses that require a combination of drugs for optimal treatment outcomes [1]. However, combining multiple medications can lead to drug interactions, which can lead to unintended side effects. These adverse effects may include reduced treatment efficacy or even drug-induced toxicity [3, 4, 5]. Such interactions, known as drug-drug interactions (DDIs), are among the leading causes of adverse drug reactions (ADRs) in patients [6]. Thus, identifying and understanding which drugs may cause DDIs is crucial. However, verifying and testing potential DDIs through conventional in vitro and in vivo experiments is costly and time consuming [6, 7].

To address these challenges, researchers have turned to alternative methods, such as in silico approaches, to predict potential DDIs [8]. With the advancement of technology, many drug-related datasets are now stored digitally. For instance, databases like DrugBank [9] compile extensive information on various drugs and make this data accessible through web interfaces and APIs. This wealth of available drug data has enabled researchers to apply computational techniques and develop models for DDI prediction, including leveraging machine learning and artificial intelligence (AI) methods [8].

Currently, there are numerous computational methods aimed at accurately predicting DDIs. In a comprehensive literature review, Han et al.[10] identified and analyzed 76 distinct DDI prediction strategies, each employing unique techniques, data sources, and drug characteristics. The development of these methods has been significantly shaped by advancements in data storage technologies, improved accessibility, and progress in statistical and machine learning models.

A large group of these approaches utilities various drug attribute similarity measures to predict DDIs with the help of machine learning models. One popular and widely used similarity measure is the SMILES (Simplified Molecular Input Line Entry System) [11] sequences of drugs, which are used in many approaches. SMILES sequences are a way of representing a molecule’s structure using a linear string of characters. This format encodes a molecule’s atoms, bonds, and connectivity in a compact, text-based form, making it useful for computational analysis and data storage. Approaches utilizing SMILES [12, 13, 14] generally exhibit favorable predictive performance. However, these approaches rely exclusively on SMILES, thereby neglecting a significant category of drugs known as biotech drugs. Biotech drugs are defined as “any medically useful drug whose manufacture involves microorganisms or substances produced by living organisms” [15]. Due to their biological nature, biotech drugs lack chemical structures and, as a result, cannot be represented by SMILES sequences. Consequently, approaches dependent on SMILES data exclude approximately 23% of the drugs listed in DrugBank [9], omitting important drugs such as penicillin and numerous vaccines.

Another challenge is the lack of reliable negative DDIs. DDI prediction approaches typically resort to labelling a random subset of unknown or unlabeled DDIs as negative for training their predictive models [16]. This method risks incorrectly classifying positive but not discovered DDIs as negative, leading to false negatives in predictions. Such misclassifications can result in models generating undesirable and potentially harmful outcomes.

To address the challenges mentioned above, we propose drug-target-protein-protein-interaction (DTPPI), a novel technique for predicting DDIs that includes biotech drugs. The DTPPI framework introduces the following key innovations:

- DTPPI constructs a weighted graph network, enabling the modeling of interactions between drugs, proteins, and targets. This network is designed to capture essential relationships regardless of whether the drugs are chemically synthesized or biotech in nature.
- DTPPI extracts six distinct topological features from the constructed graph. These features are universally applicable and can be derived for any drug type, overcoming limitations associated with chemical structure-dependent methods.
- The extracted topological features serve dual purposes: they can be used directly to predict DDIs or complement traditional drug features from databases like DrugBank, enhancing the performance of predictive models.
- DTPPI incorporates a multi-layer perceptron (MLP) prediction model, leveraging both topological and traditional features to achieve superior predictive accuracy.

Experimental results demonstrate the effectiveness of DTPPI in improving DDI prediction, including for biotech drugs typically excluded from existing methods.

## 2. Related work

Many approaches have attempted to predict DDIs with varying techniques, data preparation methods, and evaluation strategies. Similarity based approaches often rely on similarity techniques and corresponding similarity measures to predict DDIs. Gottlieb et al. [13] proposed one of these similarity-based approaches called INDI. This method calculates various similarities between drugs, constructs classification vectors for each drug, and uses pairwise combinations of these vectors as input for a logistic regression machine learning model to predict DDIs. The method utilizes seven drug attributes for similarity measures, and three target-based similarities [13]. At the time, INDI was considered a state-of-the-art similarity-based approach for DDI prediction. However, Sridhar et al. [17] argued that INDI and similar methods “neglect the structural information encoded in the biological network of drugs and their interactions”. To address this limitation, they developed a probabilistic prediction method that leverages a network of multiple drug-based similarities and known interactions. Their approach demonstrated superior predictive performance compared to INDI, as evaluated using the dataset provided by Gottlieb et al. [13]. Alternative approaches to predicting DDIs go beyond relying solely on similarity. These methods often utilize techniques such as network propagation, matrix factorization and ensemble learning. KGNN, an end-to-end framework introduced by Lin et al. [18], is designed for DDI prediction using a knowledge graph. In this graph, nodes represent various entities, such as drugs, proteins, and genes, while edges denote the relationships between them. KGNN constructs a knowledge graph incorporating multiple drugs and associated biological entities, allowing it to extract information from a drug and its neighboring nodes. This mined information is then used as features for prediction. Following KGNN, Chen et al. [12] presented MUFFIN as an approach that combines graph-based and traditional data in an ensemble framework. This method leverages the molecular structure of drugs from conventional data and the topological information of drugs from knowledge graphs [12]. Another approach is Decagon, proposed by Zitnik et al. [19]. Instead of directly predicting DDIs, Decagon focuses on identifying the specific side effects that may result from taking two drugs together. It constructs a multi-modal network that incorporates drug-drug, drug-protein, and protein-protein relationships. Features are extracted from this network and used by a convolutional neural network to predict the specific polypharmacy effects associated with pairs of drugs known to interact.

Current approaches for DDI prediction have achieved impressive results, often surpassing comparable models. However, comparisons between these methods can be misleading due to variations in data manipulation and validation metrics, and the absence of a unified benchmarking standard complicates fair evaluation. While some methods utilize graph-based techniques, biological interaction networks remain underutilized. Additionally, many approaches exclude biotech drugs, which lack chemical structures, for instance, INDI and MUFFIN rely heavily on chemical structures, making them unsuitable for biotech drugs. Moreover, some models, such as MUFFIN, use unreliable negative classes by treating unknown DDIs as negatives, which can result in false negatives.

## 3. Material and methods

### 3.1. Problem Definition

As previously stated, predicting DDIs is a problem that attempts to determine whether a pair of drugs exhibits an interaction, where the actual prediction is performed by a prediction function using a unique combination of various attributes for a pair of drugs. The DDI prediction problem can, therefore, be represented as a binary-classification problem, which can be solved using a machine-learning predictor, *f*, that approximates a function derived from the given training data, which can predict if two drugs interact 1 or not 0. *f* can be described as:

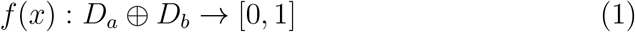

Where *D*_*a*_ and *D*_*b*_ are both 1-dimensional equally sized lists of numeric values that serve as a digital representation of drug *a* and drug *b* respectively.

### 3.2. Proposed Solution

This paper presents a novel approach for predicting DDIs, comprising three main components. The first component is a weighted DTPPI network and the motivation behind the DTPPI network can be traced back to the important role of CYP450 enzymes, which are known to significantly contribute to DDIs [20]. The DTPPI network enables the incorporation of CYP450 enzymes and other relevant proteins into a drug’s protein pathway, facilitating a more comprehensive representation of drug interactions.

The second component is the feature extractor, which vectorizes an n-dimensional numerical feature set for each drug from the DTPPI network. This feature set represents the drug based on its topology and other relevant attributes derived from the network. The feature extractor generates these topological feature sets for every drug in the DTPPI network. Additionally, the feature extractor generates traditional feature sets by referencing public data sources, such as DrugBank, for drugs not included in the DTPPI network. The final major component is the predictor which is a machine learning model that approximates a function designed to solve the DDI prediction problem. The predictor uses the combined feature set of two given drugs to make predictions, and its performance was evaluated using standard machine learning metrics.

### 3.3. Data Collection and Pre-Processing

We collected all of our data from DrugBank [9] and UniProt [21]. From DrugBank we downloaded the 2024-03-14 release of the entire database which was a 1.6 GB-sized XML-formatted data file containing all 16604 drugs, and their attributes, recorded in DrugBank. From this file we created two datasets, the first one contained 3268 unique drugs and for each drug that was extracted we included a list of drug-targets, drug-categories and a textual descriptor, if a textual descriptor was not available, the drug was not included in the dataset. Then, we extracted 720174 different DDIs based on the unique drugs found in the first dataset. From UniProt we extracted a dataset of 14853 protein-protein-interactions (PPI)s. Each pair of proteins had additional textual descriptors extracted and was labelled as either a protein or target depending on if the protein was present as a drug-target in the previous drug dataset. Table 1 presents a summary of all collected and used data for the DTPPI method.

**Table 1:** Summary of all processed data, their origin and a short description for each file.

| Dataset | Origin | Description |
| --- | --- | --- |
| DDI | DrugBank | 720174 DDIs |
| Drugs-Enriched | DrugBank | 3268 drugs with additional attributes |
| PPI | UniProt | 14853 PPIs with additional attributes |

### 3.4. Network Construction

We define the DTPPI network as an undirected weighted graph *G* =*< V, E, W >*, where *V* represents a set of nodes, which can be drugs, targets, or proteins, and *E* represents a set of edges that denote various types of interactions: drug-target interactions (DTI), target-target interactions (TTI), target-protein interactions (TPI), or PPIs and each edge in *E* is associated with a weight value. In this network, DTIs, TTIs, TPIs, and PPIs are considered homogeneous interactions, meaning there is no directionality between the interacting entities or nodes. The weight calculation for the DTPPI network was performed by computing the cosine similarity between two USE text vectors corresponding to nodes that share an edge. Cosine similarity is a measure used to assess the similarity between two non-zero n-dimensional vectors. It is defined as:

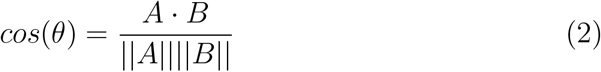

The cosine similarity value ranges from -1 to 1 and the value of 1 indicates identical vectors, 0 indicates orthogonal vectors, and -1 indicates diametrically opposed vectors. In practice, node text descriptions that are more similar to each other will receive higher edge weight values, while dissimilar text descriptions will result in lower edge weights.

### 3.5. Feature Extraction

Two sets of features were extracted using the DTPPI method: traditional and topological drug features.

- Traditional drug features are gathered from the previously mentioned collected DrugBank data. These traditional drug features are created from the “drug categories” category in DrugBank. These drug categories are short descriptive categories assigned to drugs and each drug can have multiple of these categories.
- Topological features of each drug node in the DTPPI network were analyzed using six computational methods: degree, clustering coefficient, Katz centrality, closeness centrality, degree centrality, and PageRank [22].

In the following, we defined the six informative topological features for each node in our proposed weighed mutated network.

1. The first feature extracted from the DTPPI network was the Node Degree. Node Degree describes the number of edges a given node has, and in a simple scenario, it could describe how important or popular a particular node is. The degree of a node is often used in other calculations and will be denoted as *deg*(*v*).
2. The clustering coefficient measures the extent to which the neighboring nodes of a given source node are interconnected, indicating the tightness of their connections [23]. A value of 0 means that the neighbour nodes of a given source node have no connection between each other, and a value of 1 means that the neighbour nodes are all connected. The clustering coefficient a graph can be calculated by:

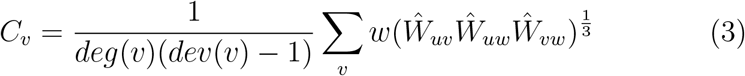
3. The Katz centrality measure is a measure of centrality that evaluates the relative influence of a node within a network. Unlike other centrality measures that may consider only direct connections, Katz’s centrality considers both the direct and indirect connections of varying lengths that a node has to other nodes in the network. The Katz centrality for node *v* can be obtained with:

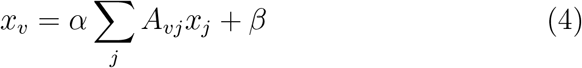 Where *A* is the adjacent matrix of the DTPPI graph with eigenvalues *λ* and parameter *β* controls the initial centrality.

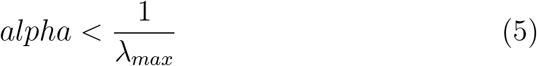
4. Closeness centrality quantifies how close a node is to all other nodes in the network [24]. It is based on the shortest paths between nodes and reflects how quickly information can spread from a given node to others in the graph as follows:

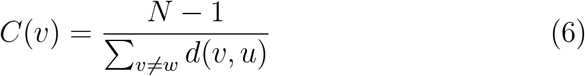
5. Degree centrality was also calculated to evaluate a node’s importance or influence through its connections to other nodes in the network. Unlike degree, it is not influenced by edge weights, therefore, it has been used as a distinct measure of connectivity [25].
6. The PageRank measures the relative importance of a particular node within a network and is defined recursively by

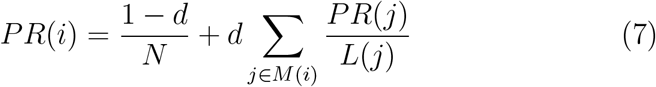

Where *PR*(*i*) is the PageRank of node *i, N* is the total number of nodes in the network, *d* is the damping factor, *M* (*i*) is the set of nodes that link to node *i* and *L*(*j*) is the number of outbound edges on node *j* [26].

## 4. Experiments

To assess the performance of the DTPPI approach, experiments were performed by training and testing a Multi-Layer Perceptron (MLP) classification model on three distinct scenarios as follows:

- **Using topological features**, in this scenario we used just the six aforementioned topological features.
- **Using biological features**, in this scenario we only used the collected drug categories as its independent features and were used as a baseline.
- **Combined features**, in this scenario we used a combination of both the topological and biological features to see if this combination has a positive influence on DDI prediction performance.

For each of these scenarios, we constructed a dataset and referred to them as the topological dataset, biological dataset, and combined dataset, respectively. Table 2 shows the detailed information of each data set.

**Table 2:** Additional information for the three topological, biological and combined datasets.

| **Data set** | **Samples** | **Features** |
| --- | --- | --- |
| Topological | 724209 | 12 |
| Biological | 724209 | 24 |
| Combined | 724209 | 36 |

These data sets were divided into a test and train data set. To mitigate potential bias arising from the substantial class imbalance between positive and negative DDI, under-sampling of the positive class in the training dataset was employed to align with the negative class ratio. The complete distribution of positive and negative classes among the data sets can be seen in Table 3.

**Table 3:** The amount of positive and negative classes per training and testing datasets.

| **Data set** | **Positive samples** | **Negative samples** |
| --- | --- | --- |
| Train | 576104 | 3263 |
| Train rebalanced | 3263 | 3263 |
| Test | 144068 | 774 |

### 4.1. Dataset pre-processing

The pre-processing began with applying MinMax scaling to all topological features to address differences in magnitudes, units, and ranges. This step ensured that no bias arose from features with higher magnitudes by scaling all values to a range between -1 and 1 while preserving their original relationships. The MinMaxScaler function from Sklearn was used for this purpose. Subsequently, biological features, initially in textual categories, were converted into numerical representations. Using Sklearn’s FeatureHasher, these features were hashed into 24-dimensional numerical vectors.

After pre-processing, the datasets were prepared for the prediction model and split into training and testing sets. 80% of the data was allocated for training, while the remaining 20% was reserved for testing.

### 4.2. Prediction Model

We used MLP as our prediction model and like any other prediction model, MLPs have specific hyperparameters that can be tuned for optimal prediction performance. To determine the optimal hyperparameters for the proposed MLP, an exhaustive grid search was performed on a cross-validation split of 5 splits. Table 4 shows the parameters used for the grid search.

**Table 4:** Parameters that been used for the exhaustive grid search cross-validation of 5 splits for the MLP prediction model

| Parameter | Values |
| --- | --- |
| solver | adam, lbfgs |
| hidden layers | (36), (36, 18), (36, 18, 9) |
| alpha | 0.1, 0.01, 0.001, 0.0001, 0.00001, 0.0000001 |
| max iter | 200, 500, 1000, 2000, 3000 |

### 4.3. Evaluation

For this work, the accuracy, precision, recall, the F1 measure and the area under the curve (AUC) of the Receiver Operating Characteristics (ROC) curve were used and reported to measure the performance of the different MLP models that were trained using either the topological, biological or combined features.

## 5. Results

### 5.1. Network

The construction of the network resulted in a connected, weighted, undirected graph of homogeneous edges and three types of heterogeneous drug, target and protein nodes. Each node in the network contained a type attribute of drug, target, or protein that was used to distinguish between the different kinds of nodes. In total, the graph consisted of 17976 unique nodes and 268307 edges between these nodes. The distribution of unique nodes consisted of 3263 drug nodes, 2886 target nodes and 11827 protein nodes. Furthermore, there were a total of 4 different edge types, namely drug-target, target-target, target-protein and protein-protein edges between the previously mentioned nodes. During the construction of the graph, 138 isolated nodes were removed due to not having any edges with other nodes. A visual representation of a subsection of the graph can be viewed in Figure 1.

**Figure 1:**
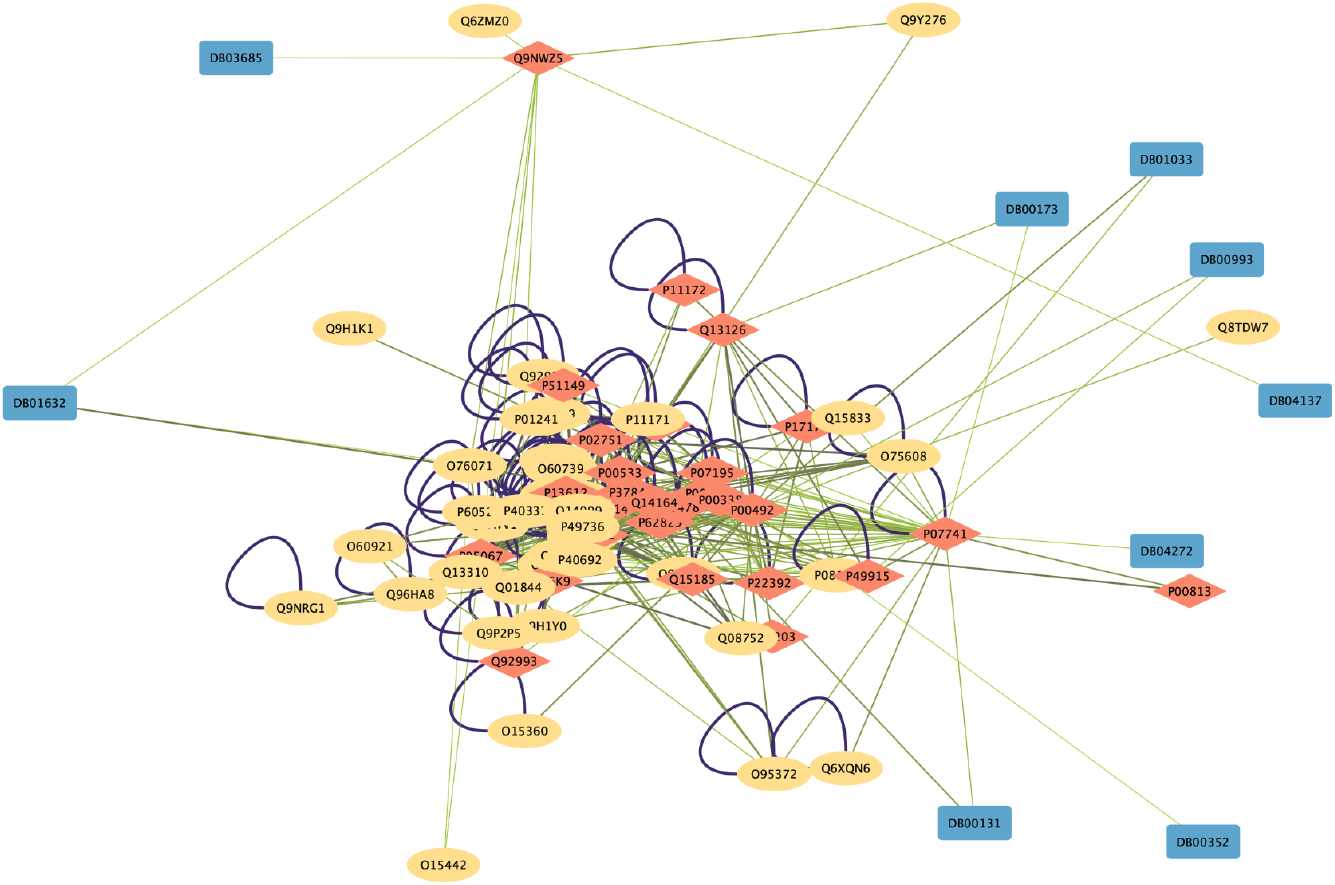
An illustration of the DB01632 drug and each two-hop neighbour node it is connected to. Blue nodes indicate drugs, red nodes target proteins and yellow nodes proteins. Each node is connected with an edge with a variant colour depending on the weight of the edge. A higher weight value results in a darker colour.

Every edge in the network contained a weight attribute ranging between 0 and 1, indicating the similarity of the USE text vectors calculated using the vectors of the two nodes connected to the edge, with 1 representing identical texts and 0 being dissimilar texts. The average edge weight within the whole network was 0.3956. The weight distribution between the four different edge types is illustrated in Figure 2 with the highest drug-target edge weight being between drug node DB09532 (Secretin human) and target node p47872 (Secretin receptor) with a weight value of 0.7818.

**Figure 2:**
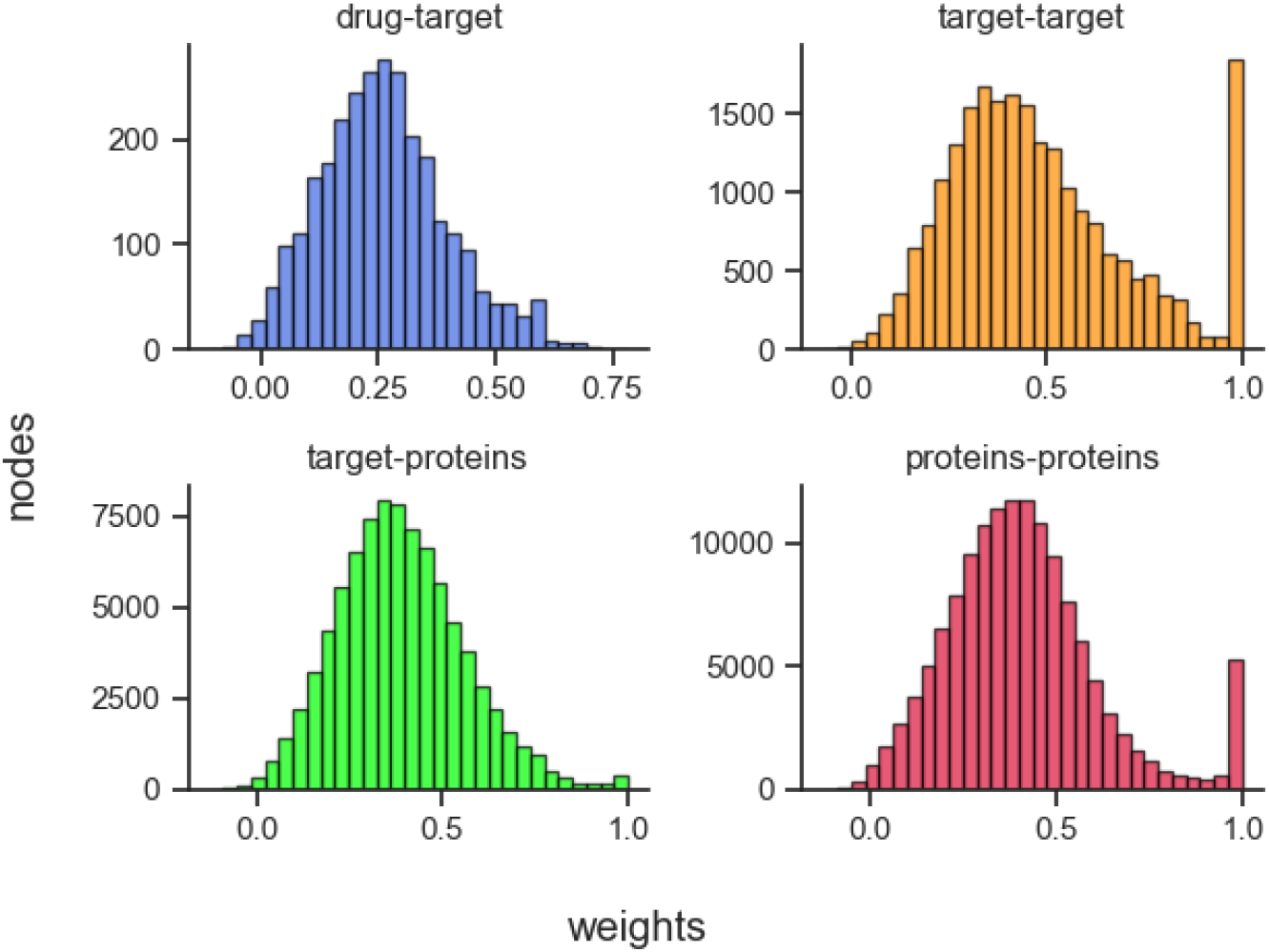
Four distribution figures with 30 bins each for every edge type found in the DTPPI network. The number of nodes each bin represents can be found along the Y-axis, and the actual weight value can be seen along the X-axis.

After the DTPPI network was constructed, two sets of the six topological features described in Section 3 in weighted and unweighted structure, were extracted. The distribution of these extracted features is illustrated in Figure 3.

**Figure 3:**
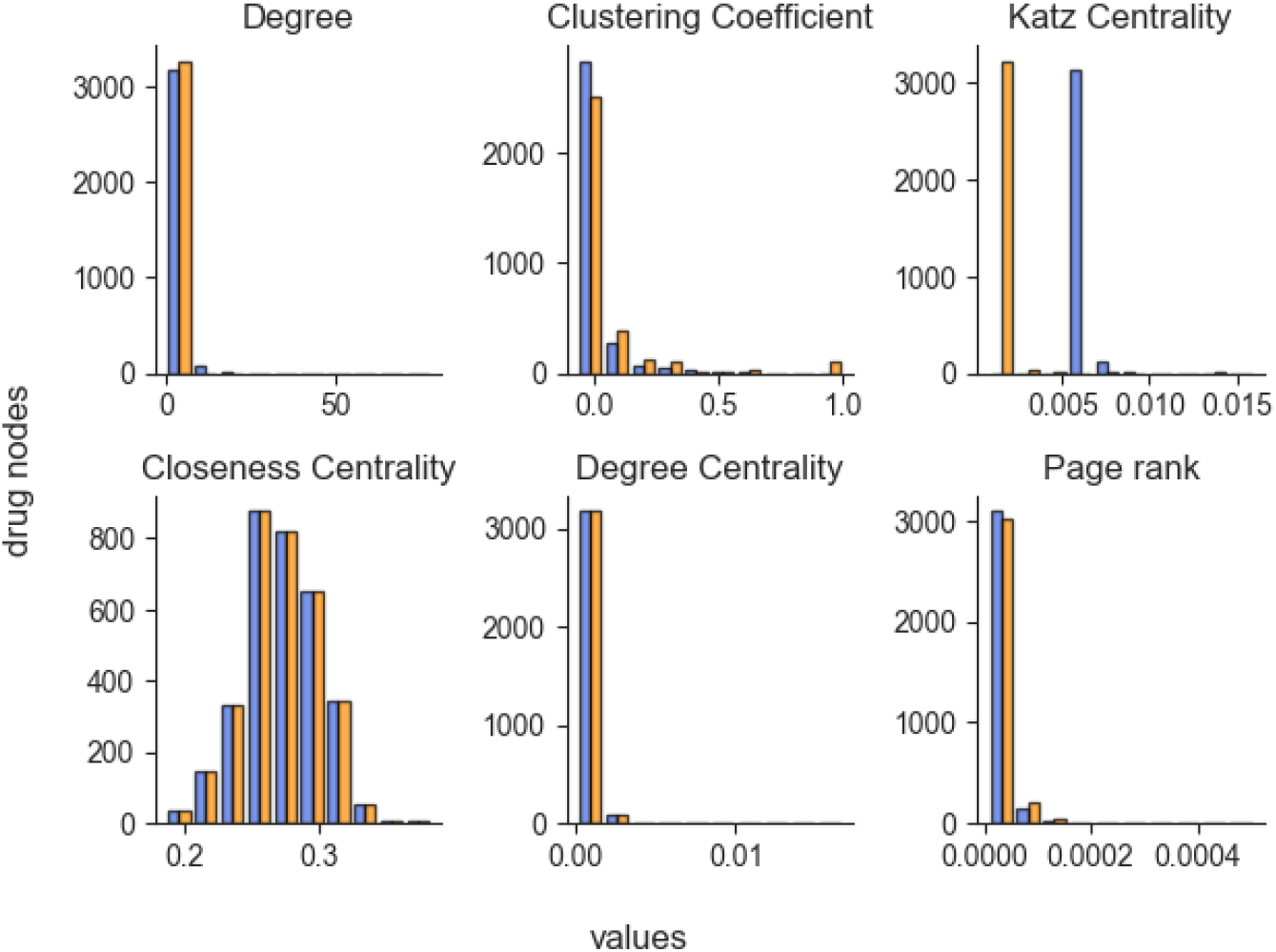
Each chart shows the distribution of values for each of the six topological values along ten bins. The number of drug nodes in each bin can be viewed on the Y-axis and the values on the X-axis. From top left to bottom right we have degree, clustering coefficient, Katz centrality, closeness centrality, degree centrality and page rank, respectively. Each chart contains weighted and unweighted values for each topological feature, with the blue bins equating weighted values and the orange bins unweighted values.

### 5.2. Model parameter fine-tuning

The prediction model was fine-tuned for optimal parameters prior to evaluation. To this end, a grid-search 5-fold cross-validation was performed for the MLP predictor for each topological, biological, and combined training dataset. Table 5 shows the result for each cross-validation per dataset. The parameters results for the biological dataset model were used for the final prediction model.

**Table 5:** MLP model variables per dataset that resulted from the 5-fold cross-validation.

| Data set | Solver | Hidden layers | Alpha | Max iter |
| --- | --- | --- | --- | --- |
| Topological | lbfgs | 36 - 18 - 9 | 1e-06 | 2000 |
| Biological | adam | 36 | 0.1 | 500 |
| Combined | adam | 36 | 0.1 | 200 |

### 5.3. Model result evaluation

The final MLP prediction model was trained and tested three times on the topological, biological, and combined datasets. For each dataset, the model was refitted and trained using the same random state to remove any possible randomness in the training of the model that would result in differing prediction results. All three prediction models were evaluated for prediction performance using the metrics discussed before. As Table 6 and Figure 4 show the complete evaluation results which confirms the combined dataset has the best performance.

**Table 6:** Resulting accuracy, precision, recall, f1 and AUC evaluation per train test phase for the three topological, biological and combined datasets.

| Phase | Data set | Accuracy | Precision | Recall | F1 | AUC |
| --- | --- | --- | --- | --- | --- | --- |
| train | Topological | 0.5932 | 0.6466 | 0.4110 | 0.5026 | - |
|  | Biological | 0.9012 | 0.8936 | 0.9108 | 0.9021 | - |
|  | Combined | 0.9186 | 0.9152 | 0.9228 | 0.9190 | - |
| test | Topological | 0.4099 | 0.9970 | 0.4080 | 0.5790 | 0.6441 |
|  | Biological | 0.8128 | 0.9987 | 0.8129 | 0.8963 | 0.8920 |
|  | Combined | 0.8258 | 0.9989 | 0.8258 | 0.9041 | 0.9143 |

**Figure 4:**
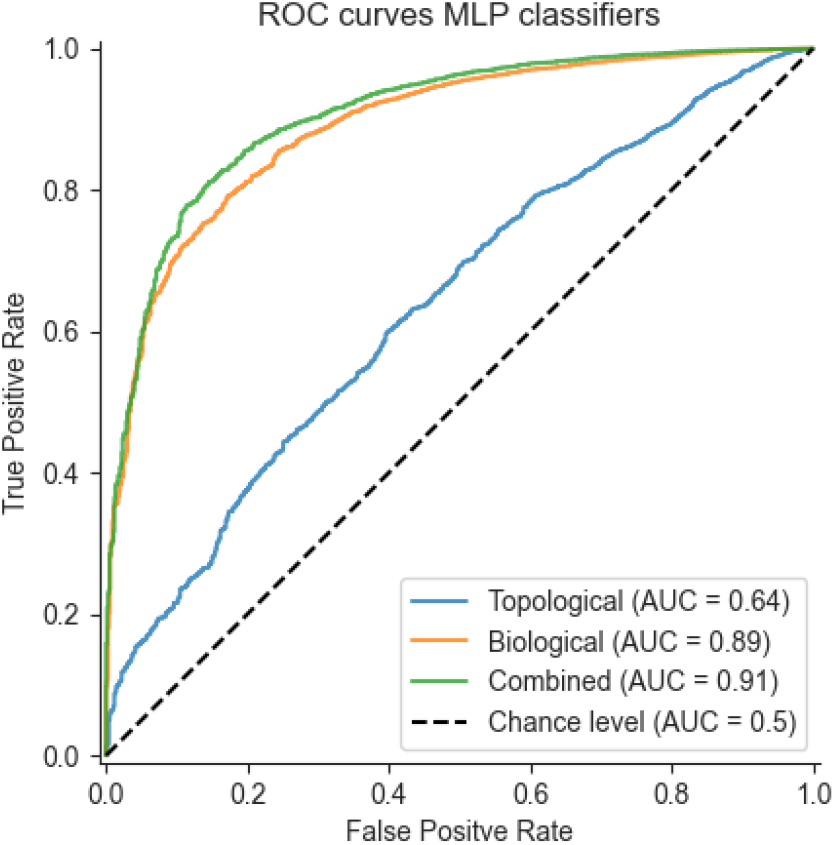
This Figure represents the ROC curve for the three topological, biological and combined datasets. It also includes a baseline “chance” level ROC curve equating to a random prediction model.

Figure 4 shows an AUC performance of 0.64 for the topological-trained model, 0.89 for the model trained on biological features and an AUC of 0.91 for the final model trained on the combination of topological and biological features.

## 6. Discussion

The main goal of this paper was to research the viability of a novel approach for predicting DDIs that, in some way, leverages information-rich biological networks. The results show that an increase of AUC, and all other DDI prediction evaluation metrics, can be expected of a predictor model trained on a dataset consisting of more traditional biological features reinforced with topological features originating from the DTPPI network. It is important to note that topological features alone were insufficient to achieve acceptable DDI prediction performance metrics.

Whilst predictive performance using both combined topological and biological features is good, it is essential to note the imbalance of positive DDI classes compared to negative classes. The training data was undersampled to an even class distribution to combat this imbalance. However, the overwhelming imbalance causes unreliable precision metric results, with near-perfect scores for all test dataset variations; these test datasets did not undergo the under-sampling mentioned above. However, the biological and topological feature sets still outperformed other evaluation metrics. A more substantial amount of generated or verified negative classes would result in a better generalised and realistic prediction model.

Additional findings suggest that using weighted topological features from a biological network is a viable mechanism for DDI prediction approaches as an alternative to other, often more complex and computationally heavy, network-based approaches that make use of unweighted networks such as [18, 12]. This difference between having a weighted and unweighted network is mainly supported by the results in Figure 2, which shows a different feature relationship, primarily in the Katz Centrality and Clustering Coefficient. Therefore, the usage of USE text embedding as weighting mechanics, which up until this point has not yet been utilised in other related research, is a viable approach, especially when a similarity measure is required between a drug and a biological entity such as a protein. It must be noted that USE is not made to embed large and complex biological texts. A more fine-tuned text embedding approach, tuned explicitly for complex biological and pharmacological texts, would result in even more accurate, varied, and informative features for weighted networks than their unweighted counterparts.

The DTPPI approach has shown that simple graph topological features increase DDI predictive performance. Further research is recommended regarding more advanced graph feature extraction techniques, such as node representation learning or even techniques such as graph neural networks. These techniques can lead to even greater predictive performance without altering the DTPPI network.

Additionally, the usage of reliable negative DDI classes has been shown to affect the generalisability of this research. Further research is recommended to increase the amount of reliable negative DDI data. This will allow other related DDI approaches that model the DDI prediction problem as a binary classification problem to have an increased overall dataset, which will help with the generalisability and predictive performance of said approaches, including DTPPI.

## References

[1] A. Calderón-Larrañaga, B. Poblador-Plou, F. González-Rubio, et al., Multimorbidity, polypharmacy, referrals, and adverse drug events: are we doing things well?, British Journal of General Practice 62 (605) (2012) e821–e826.

[2] G. Taheri, M. Habibi, T. Sedghamiz, Machine learning-based prediction for drug-drug interaction using a knowledge graph.

[3] W. Y. Zheng, L. C. Richardson, L. Li, R. O. Day, J. I. Westbrook, M. T. Baysari, Drug-drug interactions and their harmful effects in hospitalised patients: a systematic review and meta-analysis, European Journal of Clinical Pharmacology 74 (1) (2018) 15–27.

[4] M. Ayati, G. Taheri, S. Arab, L. Wong, C. Eslahchi, Overcoming drug resistance by co-targeting, in: 2010 IEEE International Conference on Bioinformatics and Biomedicine (BIBM), 2010, pp. 198–202.

[5] G. Taheri, M. Ayati, L. Wong, C. Eslahchi, Two scenarios for overcoming drug resistance by co–targeting, International journal of bioinformatics research and applications 11 (1) (2015) 72–89.

[6] C. Palleria, A. Di Paolo, C.e.a. Giofré, Pharmacokinetic drug-drug interaction and their implication in clinical management, Journal of Research in Medical Sciences : The Official Journal of Isfahan University of Medical Sciences 18 (7) (2013) 601–610.

[7] G. Taheri, M. Habibi, Identification of essential genes associated with sars-cov-2 infection as potential drug target candidates with machine learning algorithms, Scientific Reports 13 (1) (2023) 15141.

[8] R. Safdari, R. Ferdousi, K. Aziziheris, S. R. Niakan-Kalhori, Y. Omidi, Computerized techniques pave the way for drug-drug interaction prediction and interpretation, BioImpacts: BI 6 (2) (2016) 71–78.

[9] D. S. Wishart, Y. D. Feunang, A. C. Guo, E. J. Lo, e. a. Marcu, Drug-Bank 5.0: a major update to the DrugBank database for 2018, Nucleic Acids Research 46 (D1) (2018) D1074–D1082.

[10] K. Han, P. Cao, Y. Wang, F. e. a. Xie, A Review of Approaches for Predicting Drug–Drug Interactions Based on Machine Learning, Frontiers in Pharmacology 12 (2022).

[11] D. Weininger, SMILES, a chemical language and information system. 1. Introduction to methodology and encoding rules (2002).

[12] Y. Chen, T. Ma, X. Yang, J. Wang, B. Song, X. Zeng, MUFFIN: multiscale feature fusion for drug–drug interaction prediction, Bioinformatics 37 (17) (2021) 2651–2658.

[13] A. Gottlieb, G. Y. Stein, Y. Oron, E. Ruppin, R. Sharan, INDI: a computational framework for inferring drug interactions and their associated recommendations, Molecular Systems Biology 8 (1) (2012) 592.

[14] B. Ran, L. Chen, M. Li, Y. Han, Q. Dai, Drug-Drug Interactions Prediction Using Fingerprint Only, Computational and Mathematical Methods in Medicine 2022 (2022) e7818480.

[15] F. M. Steinberg, J. Raso, Biotech pharmaceuticals and biotherapy: an overview, Journal of Pharmacy & Pharmaceutical Sciences: A Publication of the Canadian Society for Pharmaceutical Sciences, Societe Canadienne Des Sciences Pharmaceutiques 1 (2) (1998) 48–59.

[16] Y. Zheng, H. Peng, X. Zhang, Z. Zhao, X. Gao, J. Li, DDI-PULearn: a positive-unlabeled learning method for large-scale prediction of drugdrug interactions, BMC Bioinformatics 20 (19) (2019) 661.

[17] D. Sridhar, S. Fakhraei, L. Getoor, A probabilistic approach for collective similarity-based drug–drug interaction prediction, Bioinformatics 32 (20) (2016) 3175–3182.

[18] X. Lin, Z. Quan, Z.-J. Wang, T. Ma, X. Zeng, KGNN: Knowledge Graph Neural Network for Drug-Drug Interaction Prediction, in: Proceedings of the Twenty-Ninth International Joint Conference on Artificial Intelligence, Yokohama, Japan, 2020, pp. 2739–2745.

[19] M. Zitnik, M. Agrawal, J. Leskovec, Modeling polypharmacy side effects with graph convolutional networks, Bioinformatics 34 (13) (2018) i457–i466.

[20] P. Manikandan, S. Nagini, Cytochrome P450 Structure, Function and Clinical Significance: A Review, Current Drug Targets 19 (1) (2018) 38–54.

[21] The UniProt Consortium, UniProt: the Universal Protein Knowledge-base in 2023, Nucleic Acids Research 51 (2023) D523–D531.

[22] S. Brin, L. Page, The anatomy of a large-scale hypertextual Web search engine, Computer Networks and ISDN Systems 30 (1) (1998) 107–117.

[23] W. L. Hamilton, Graph Representation learning, Vol. 14, Morgan and Claypool, 2020.

[24] M. Habibi, G. Taheri, A new machine learning method for cancer mutation analysis, PLoS computational biology 18 (10) (2022) e1010332.

[25] M. Habibi, G. Taheri, Topological network based drug repurposing for coronavirus 2019, PLOS ONE 16 (7) (2021) e0255270.

[26] G. Taheri, M. Habibi, Uncovering driver genes in breast cancer through an innovative machine learning mutational analysis method, Computers in Biology and Medicine 171 (2024) 108234.

